# Effectiveness of surface-based North Atlantic right whale detection methods for vessel strike mitigation

**DOI:** 10.1101/2021.08.27.457997

**Authors:** Loïcka M.R. Baille, Daniel P. Zitterbart

## Abstract

Increasing commercial and recreational use of the world’s ocean leads to growing concerns on vessel and marine mammal encounters. For endangered species, like the North Atlantic right whale (NARW), vessel strikes can be responsible for the majority of the recorded deaths. Reducing the number of vessel strikes is key to improve North Atlantic right whale protection and a number of mitigation methods have been proposed and implemented. In this manuscript, we developed an agent-based model to assess the effectiveness of surface-based whale detection methods for vessel strike mitigation. We find that the effectiveness of such systems varies highly depending on the vessel’s speed and maneuverability. We also find that if vessel-based whale detection systems are used in conjunction with other mitigation measures such as general speed restrictions, they can be very effective and could lead to a significant decrease in vessel strikes when deployed at a large-scale.

## 1 INTRODUCTION

### 1.1 Vessel strikes worldwide

Cetaceans face a multitude of anthropogenic threats, such as vessel strikes, ocean pollution, ocean noise, climate change, fishing gear entanglement, and even whaling in some parts of the world (Sébe et al. 2019). A vessel or ship strike is defined as any physical impact, fatal or not, between any part of a watercraft and a live marine animal (Peel et al. 2018). Vessel strikes and entanglement have become main concerns as the world’s oceans have been experiencing an increasing level of use due to growing commercial and recreational activities (Sébe et al. 2019). In 1890, the total recorded world fleet amounted to 11,108 commercial vessels (>100 gross tons), while in 2020, over 98,000 were accounted for, which is equivalent to a 783% growth (Laist et al. 2001, NATIONS n.d.). This surge in maritime traffic led to growing concerns as vessel strikes do impact marine life welfare, crew’s safety, and lead to negative economic consequences (Schoeman et al. 2020). With the growing interest in autonomous vessels, even greater numbers of vessels are expected to travel the oceans, hence increasing the risk of vessel strikes. Since 2005 vessel strikes have been identified as a priority by the International Whaling Commission Conservation Committee (IWC-CC). IWC-CC aims to identify at-risk populations and high-risk areas to develop and implement solutions to achieve a permanent reduction in vessel strikes (International Whaling Commission). Schoeman et al. (2020) established that at least 75 marine species are being affected by vessel strikes, from North Atlantic Right Whales (NARWs), dolphins, to penguins, and fish.

### 1.2 Vessel strikes NARW

The North Atlantic right whale (NARW, *Eubalaena glacialis*) is one of the world’s most endangered large whale species, with approximately 360 individuals remaining in 2019 (Moore et al. 2021). Fishing gear entanglement and vessel strikes are responsible for at least 86 mortalities and serious injuries in the US and Canada between 2000 and 2017 (NOAA Fisheries, Office of Protected Resources 2020). Due to low numbers, high mortality rates and low calving rates the species has been classified as “Endangered” under the Endangered Species Act since 1970. While most large whale species are vulnerable to vessel traffic (Laist et al. 2001), NARWs are two orders of magnitude more prone to vessel strikes (Vanderlaan & Taggart 2007). This is due to their surface-skimming feeding behavior, and their habitats and migration routes often overlapping with ports and/or shipping lanes (Parks et al. 2012, Fisheries Thu, 06/03/2021 - 11:54). Vessel strikes account for 52.5% of all deaths among necropsied right whales between 1970 and 2006 (Campbell-Malone et al. 2008). Assessing the true number of NARW vessel strikes is still a difficult task, as striked animals might not be stranded or found. Pace et al. (2021) estimated that only 36% of all estimated NARW deaths were accounted for via observed carcasses between 1990 and 2017 (Pace et al. 2021). Estimates of annually striked animals vary from 0.81 (Hayes et al. 2018) to tens or hundreds (Conn & Silber 2013) whales per year.

### 1.3 Large Scale vessel strikes mitigation strategies

During the last decades a variety of mitigation measures have been developed to reduce the probability of vessel strikes. Vessel strike mitigation approaches can be classified into either large-scale approaches, where high-risks areas are established and a set of navigation rules are implemented, and small-scale approaches, that rely on vessels to detect at-risk animals and subsequently alter their course and speed to avoid collision. It is to be noted that marine mammal observations from vessel-based methods can always be used to inform large-scale mitigation efforts about the presence of an animal at a given location and time. In the following, we will provide a brief overview of the existing mitigation efforts for North Atlantic right whales.

#### 1.3.1 Re-routing

Rerouting aims to separate vessels from areas that are highly used by NARWs. Once high-risk regions are established, alternative vessel routes may be created to avoid such areas. Routing measures may be permanent, seasonal, mandatory or recommended, and may apply to all vessels or only a subset (Schoeman et al. 2020). Currently the following implementations of re-routing are used:

##### Area To Be Avoided (ATBA)

ATBAs discourage vessels from transiting through certain areas of the ocean. Specifically, the International Maritime Organization (IMO) defines an ATBA as “a routeing measure comprising an area within defined limits in which either navigation is particularly hazardous or it is exceptionally important to avoid casualties and which should be avoided by all vessels, or certain classes of vessels” (*Ships’ Routeing* n.d.). Seasonal ATBAs in the Great South Channel, Cape Cod Bay, MA and Roseway Basin, Canada were established to protect North Atlantic Right Whales (Vanderlaan & Taggart 2009).

##### Dynamic Management Areas (DMAs)

In the US, voluntary DMAs, also called Slow Zones, are discrete areas established by NOAA Fisheries, where visual sightings of three or more North Atlantic right whales have been recorded within 15 days (Fisheries Thu, 06/03/2021 - 12:38). Mariners are encouraged to avoid these temporary areas or reduce their speed to 10 knots or less when transiting through them to avoid vessel strike (Fisheries Thu, 06/03/2021 - 12:38). NOAA created the Right Whale Sightings Advisory System to collect, validate, and communicate visual sightings reported by individuals. Those reports may also be used to establish new DMAs (Johnson et al. 2020).

##### Seasonal Management Areas (SMAs)

When passing through SMAs, all vessels 65 feet (19.8 meters) or longer must travel at 10 knots or less, along the U.S. east coast (Cape Cod Bay, off Race Point, Great South Channel), at certain times of the year to reduce the threat of vessel collisions with endangered North Atlantic right whales (Services n.d.). Van der Hoop et al. (2015) observed that large whale mortalities due to vessel strikes decreased when SMAs are active compared to when SMAs are not active (van der Hoop et al. 2015).

##### Traffic Separation Schemes (TSSs)

TSSs are routing measures that separate opposing streams of traffic through the establishment of traffic lanes. Certain of these permanent mandatory routes have been amended to reduce the co-occurrence of vessels and whales (Santa Barbara Channel, San Francisco Bay, Bay of Fundy (Canada), Boston) (Fisheries Thu, 06/03/2021 - 12:38, Schoeman et al. 2020).

#### 1.3.2 Speed restrictions

It has been shown that reducing vessels’ speed decreases vessel strikes’ rate and the injuries’ severity (Vanderlaan & Taggart 2007, Gende et al. 2011, Conn & Silber 2013). Laist et al. (2001), and later Conn and Silber (2013), showed that the probability of lethal injury decreased to lower than 50% when traveling at speeds *leq*10 knots. In both studies, the vessel strike rate also decreased for lower vessel speeds. Proposals for speed restrictions can be submitted to the International Maritime Organization to implement, voluntary or mandatory, permanent or seasonal, vessel speed restriction zones outside of territorial waters (Silber, Vanderlaan, Tejedor Arceredillo, Johnson, Taggart, Brown, Bettridge & Sagarminaga 2012). A reduction in vessel speed is the preferred measure to implement when vessels cannot be re-routed (Schoeman et al. 2020).

### 1.4 Small Scale vessel strike mitigation strategies

The desired vessel strike mitigation measure would be applied on an individual vessel basis, where each transiting vessel would be in charge of detecting at-risk animals and react accordingly, by slowing down and changing its course to minimize the risk of vessel strike (Weinrich et al. 2010, Flynn & Calambokidis 2019). Such mitigation measures can be implemented on-top of large-scale mitigation strategies or independently (Wiley et al. 2016).

### 1.5 Detection Methods

In order to establish high-risk areas and/or to implement small-scale mitigation measures, a multitude of methods have been developed to detect large marine mammals.

#### 1.5.1 Passive Acoustic Mitigation (PAM)

PAM capabilities have massively improved over the last two decades as acoustic monitoring overcomes some of the limitations visual monitoring faces, such as bad weather (Verfuss et al. 2018). PAM relies on underwater microphones, hydrophones, to detect, classify and/or localise marine mammals’ vocalizations, from a few hundred meters away to several kilometers depending on the environmental condition and species’ vocalization frequencies (Verfuss et al. 2018). Hydrophones can either be permanently moored down or towed by a vessel, more commonly on seismic surveys (Verfuss et al. 2018). Moored PAM systems, in the form of auto-detection buoys with hydrophones located 60-120ft below the surface, have been implemented in the Port of Boston, Cape Cod Bay, the coasts of Georgia and Florida (Baumgartner et al. 2019). These PAM systems constantly listen for NARW calls and send potential detections to a command-and-control center where trained analysts validate the sound. If the call is within 5-nautical miles of the buoy, alerts are sent out via radio, email, online (Right Whale Listening Network) to LNG tankers with re-routing or speed reduction instructions (Knowlton 2020). However, PAM relies on animals to vocalise frequently and on background noise to be low enough to not interfere with vocalizations (Zitterbart et al. 2020). Hence, this method can lead to varying results due to environmental conditions, equipment, deployment types and target species (Verfuss et al. 2018). Very few of those systems are suitable for small-scale vessel strike mitigation approaches due to the logistical effort of towing hydrophone systems capable of detecting animals vocalizing in front of the vessel.

#### 1.5.2 Marine Mammal Observers (MMOs)

Manual detection of marine mammals via dedicated observers is still the most prevalent method used for any mitigation purposes (Weinrich et al. 2010, Zitterbart et al. 2020). Trained marine mammal observers scan the ocean surface surrounding the vessel, up to 5000 m, for potential sightings (Pyc et al. 2015). Weunrich et al. (2010) showed that MMOs are more likely to detect animals than other crew members thanks to their experience and their lack of distractions from other factors. However, marine mammal observers are impacted by weather conditions and can only work at night in conjunction with night vision goggles, which greatly reduces the effectiveness (Schoeman et al. 2020). Weinrich et al. (2010) found that trained marine observers significantly increased the number of sightings on highspeed vessels and effectively prevented vessel strikes. During their study’s time frame, the ferry using a dedicated MMO did not experience any strikes, while a similar boat without MMOs, transiting through the same route, collided with a fin whale (Weinrich et al. 2010).

#### 1.5.3 Active Acoustic Monitoring (AAM)

Active acoustics has successfully been implemented to detect marine mammals up to 2000 m in front of a vessel via active sonars (Pyc et al. 2015). This technique emits pulses of sounds and records returning echoes to localize objects. Active acoustic methods inadvertently increase noise levels in the water, which can be detrimental to marine species (André et al. 2011) and performance highly varies with the prevalent sound propagation conditions.

#### 1.5.4 Radio Detection and Ranging (RADAR)

RADAR systems emit electromagnetic waves, and record for returning echoes to determine size, shape, distance, and speed of a target. RADAR technology aims to detect surface targets such as an animal body part, exhalations or sea surface disturbances (Verfuss et al. 2018). It operates best at detection ranges under a kilometer and at low sea state conditions as the shorter wavelength of electromagnetic waves are rapidly absorbed by water molecules (Verfuss et al. 2018).

#### 1.5.5 Thermal Infrared Imaging (Thermal IR)

Thermal IR scanners are passive imaging systems that can be used day and night, on land (Zitterbart et al. 2020) and vessels (Zitterbart et al. 2013). They rely on an apparent temperature difference between the above-surface body parts of the animal or its exhalation and the ocean. Thermal IR systems have been shown to detect large whales reliably up to several kilometers away (Zitterbart et al. 2020).

The aim of this study is to assess the effectiveness of surface detection methods (i.e. detecting the animal when it is at the surface), such as thermal imaging, radar or marine mammal observers for vessel strike mitigation of NARWs. This was achieved by creating an agent-based model, where vessel strike risk can be assessed for different vessels and animal characteristics. The agents’ behavior was derived from experimentally collected NARW dive profiles (Baumgartner & Mate 2003, Baumgartner et al. 2017). We find that the detection performance and the vessel characteristics (speed, and capacity to change vessel’s course and speed) have the highest impact on the ability to detect a NARW early enough to still take evasive action. Furthermore, we find that when vessel-based mitigation strategies are paired with large-scale mitigation approaches, such as speed restriction (10kn), significant levels of protection can be achieved.

## 2 MATERIALS AND METHODS

### 2.1 WHorld (the grid)

A 3-dimensional grid was generated and virtual vessels and whales (animats) were distributed randomly on that grid and instructed to move for 60min (in model time). Each vessel was placed on the grid with a fixed speed and trajectory. Similarly, each animat was placed on the grid and instructed to move in the horizontal plane according to a correlated random walk with a random +/-5° change in heading between steps, a fixed speed, and a uniquely generated dive profile. The online supplementary material provided along this manuscript summarizes the sets of parameters used for each of the simulated cases.

### 2.2 Whales

#### 2.2.1 Dive profiles

Each animat followed an artificially generated dive profile. To mimic true NARW dive behavior, we extracted the diving characteristics from biologging data collected in the summers of 2000 and 2001 via suction-cup mounted time-depth recorders (TDR) (Baumgartner & Mate 2003, Baumgartner et al. 2017). Using those characteristics, a unique dive profile was generated for each animat. Artificial dive profiles were generated as follows: time-depth data from TDR were manually selected and trimmed for quality control, e.g. remove time sections when the tag fell off the animal. Dive profiles were classified by a human analyst as either shallow or deep dives, depending on the whales feeding behavior inferred from the dive profile. Subsequently, time-depth data were segmented into three depth layers: surface [0 - 5m], subsurface [5 - 10m], and deep 10+m, as those sections determine the whale’s availability bias as well as their vulnerability to vessel strikes. We used the distributions of duration, occurrences, and number of transitions between depth sections to generate artificial dive profiles. For simulation purposes, four behavioral states were established. State 0 corresponds to the whale blowing at the water surface, making the animat available for detection. State 1 and state 2 correspond to a whale located 0-5m and 5-10m deep, respectively. In both states 1 and 2, the whale is considered vulnerable to vessel strikes as the hull of many vessels can reach such depths,making vessel strikes with the diving animal possible. Finally, when a whale dives deeper than 10m, it is considered to be out of reach of most non-container vessels’ hulls and therefore not susceptible to vessel strikes. Larger container vessels that can have drafts up to 15.3m (Ultra Large Container Vessel) are not considered in this study because we assume their maneuverability is so limited that effective mitigation measures through evasive maneuvers could not be implemented (Wikipedia 2021).

During the simulation, we studied three different animat behaviors for shallow, deep and mixed diving behavior. Shallow and deep artificial dive profiles were exclusively generated from dive profiles classified as shallow and deep respectively, while in mixed behavior, all dive profiles were considered.

#### 2.2.2 Inter-blow Interval (IBI)

We define an animal as available for detection when it is exhaling at the surface, because that is the main cue used by thermal imaging detection systems and marine mammal observers when the animal is far enough away for evasive actions to still be feasible (Zitterbart et al. 2013). Times of exhalation cannot be extracted from dive profiles. We therefore incorporated exhalations (state 0, 3 seconds long) into the artificial dive profiles while the animat is at the surface (state 1) at certain intervals, defined as the inter-blow interval (IBI). In this simulation, 30, 60, 120,and 300sec IBIs were tested.

### 2.3 Vessels

Our aim for this simulation was to provide an assessment for a broad range of vessels. Maritime vessels that are susceptible to vessel strikes come in a large variety of vessel configurations, many more than we could consider within the scope of this study. The same is true for the variety of detection systems. Therefore, we chose the vessels and detection systems parameters to be broad enough so that they apply to a wide range of vessel categories. A summary of the vessel’s parameters used is provided in Table 1. We modeled vessel speeds from 1-15m/s to account for a wide range of vessels, from fishing vessels to high-speed ferries. The field of view of the detection system was chosen to be 20°, which is a reasonable assumption because the vessels’ speed is usually several times higher than a whale’s swimming speed, therefore it is highly unlikely for a whale to enter a vessel’s path from beyond that area. Each vessel was modeled with a specific and constant reliable detection range. The reliable detection range (RDR) is defined as the maximum distance at which a whale blow would be detected with certainty (probability of detection (PD) = 1). To account for a wide variety of vessel types, we define a ship’s Reaction Time (RT) as the time a vessel requires to make an effective mitigation maneuver. This variable integrates covariates such as a vessel’s maneuverability, its ability to slow down, and the time needed between detection and alert.

**Table 1:**
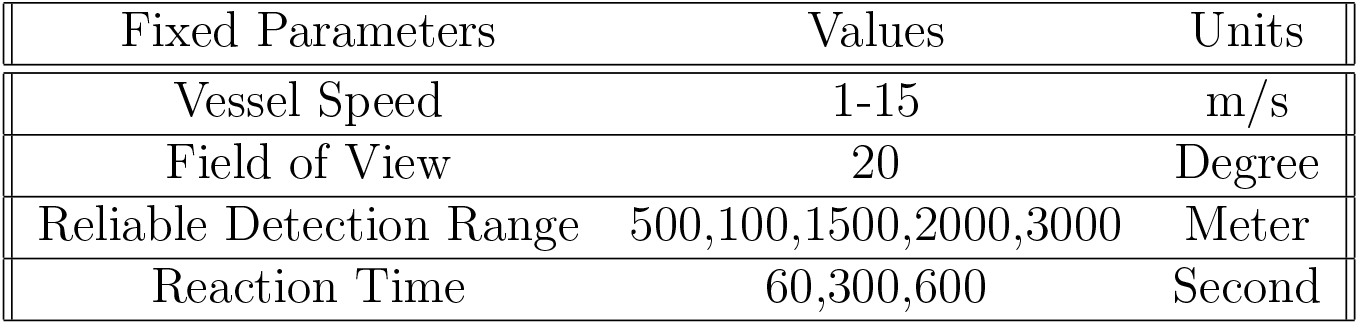
Vessel parameters used for agent-based simulation

### 2.4 Detection

### 2.4.1 Detection Function

We used a detection function, defined as the probability to detect a whale’s blow at a given distance, obtained during a previous experiment (Zitterbart et al. 2020). Humpback whale blow detection data was collected using a thermal IR imaging camera off Poipu Shores, Hawaii in 2016, comparable to a point-transect distance sampling detection scheme. We derived the detection function by fitting a log-normal distribution to the detection data. The range at which the fitted detection function peaks (1600m) was used as the furthest distance a whale blow would be detected with certainty (e.g. probability of detection (PD) = 1), previously defined as Reliable Detection Range (RDR).

To assess the impact of the shape of the detection function, i.e. detections beyond RDR (PD¡1), we tested two different scenarios. In the RDR scenario, the detection probability is binary, set to 1 for ranges below or equal to RDR, and 0 for ranges beyond RDR. In the Data-driven Detection Function (DDF) scenario, the detection probability is set to 1 for ranges below or equal to RDR, and logarithmically decreases according to the detection function (Figure 3 A).

To test different reliable detection ranges, either in the RDR or DDF scenarios, the detection function was simply moved along the x-axis, while the shape of the detection function was kept similar (Appendix **??**).

#### 2.4.2 Detection performance metric

For successful mitigation, a whale has to be detected early enough (in-time) so the vessel can still take evasive actions (Zitterbart et al. 2013). This can be parameterized with the definition of a safe zone and a danger zone (Figure 1). We define the danger zone as the area where if a whale was detected there, the vessels would be too close to safely maneuver to avoid a strike. The danger zone range (DZR) is derived from the vessel’s speed and the reaction time (DZR=RT*vessel speed). On the other hand, the safe zone represents the area from the DZR to the far end of the detection range. When a whale is detected in this zone, vessels have enough time to implement the necessary measures to safely alter their trajectories and avoid a strike. In our study, a whale is detected in-time when it was spotted in the safe zone, before it entered the danger zone. We define the In-Time Detection Probability (ITDP) as our metric to evaluate the impact of the different vessel and whale parameters. Only whales that would enter the danger zone are considered. The ITDP is defined as the proportion of whales that were detected in the safe zone from all whales that entered the danger zone.

**Figure 1:**
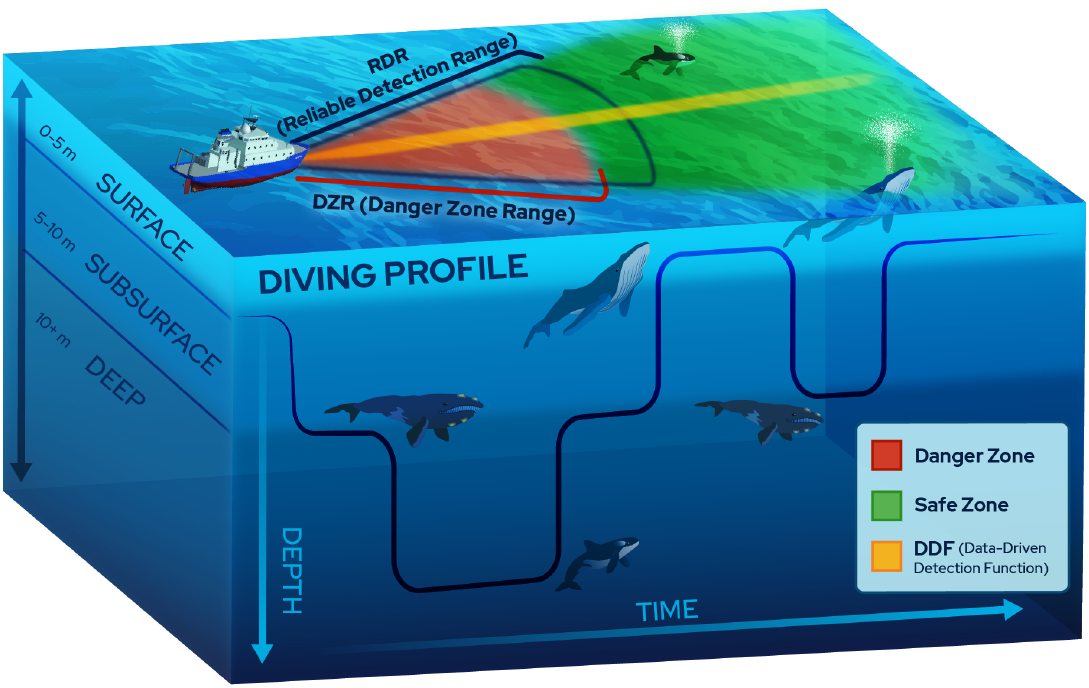
Definitions of relevant areas and dive states. The detection area of a surface-based detection system with a 20 degree field of view is divided into a danger zone and a safe zone, which depend on vessel speed, vessel reaction time, and the Reliable Detection Range (RDR) of the mitigation system. For the purpose of this study, the whales’ possible depths were divided into three categories: surface [0-5m], subsurface [5-10m] and deep [10+m]. Only whales that are at the surface and blowing can be detected by a vessel

**Figure 2:**
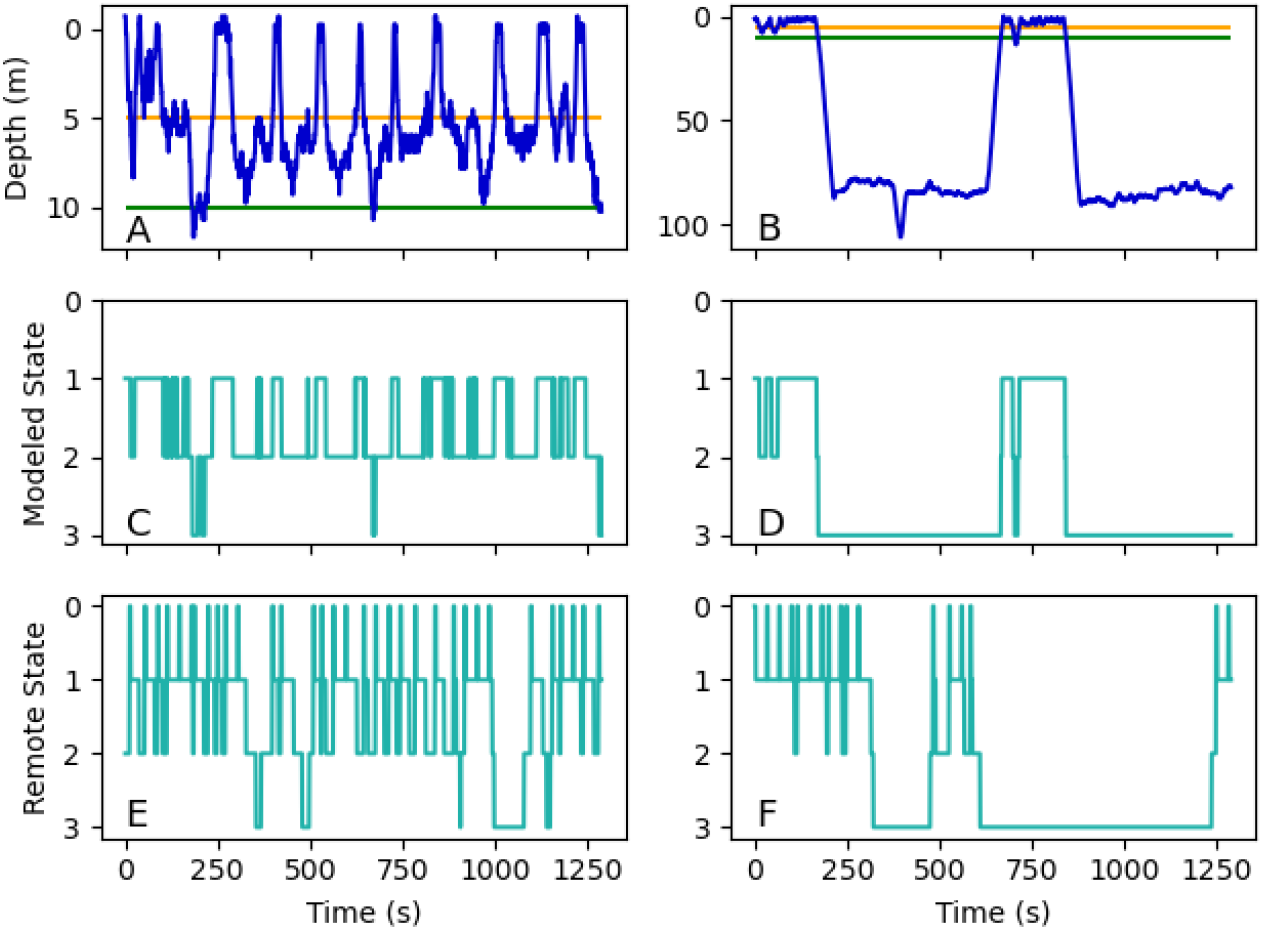
Artificial dive profiles are generated using a three-steps process. The left column (A,C,E) depicts the process when applied to a shallow dive behavior, while the right column (B,D,F) is applied to a deep dive behavior. The first row (A,B) illustrates true time-depth data collected via TDRs. The horizontal 5m (orange) and 10m (green) lines mark the boundaries between states 1, 2 and 3. State 1 includes depths between 0 and 5m, stage 2 encompasses depths from 5 to 10m, and stage 3 is comprised of 10m+ depths. The second row (C,D) shows the conversion from depth values to state values (0,1,2,3) based on the respective true dive profiles from the first row. Finally, the third row (E,F) shows possible examples of modeled state dive profiles derived from the respective distributions of duration, occurrences, and number of transitions between depth sections of the true time-depth data. State 0 refers to whale exhalations that were artificially added at set intervals (inter-blow intervals) since collected time-depth data did not provide such information.

The simulation code and supporting data can be found in this repository: https://github.com/whoi-mars/WHorld_public.git

## 3 RESULTS

### 3.1 Detection Function Shape

We find that the shape of the detection function (Figure 3B; Appendix **??**) has a significant impact on the ITDP. When we do not consider whales that are detected at distances where the probability of detection is less than 1 (RDR scenario, 1000m, Figure 4A), the ITDP drops to values below 90% for vessel-reaction time (RTs) above 1.5min, and to zero for RTs above 3min (Figure 4A). This can be explained by the linear increase of the danger zone with reaction time, for a given speed (5m/s in this example). Due to a fixed detection range, a larger danger zone reduces the safe zone and vice versa. Beyond a certain RT, the danger zone range will be larger than the reliable detection range, thus the ITDP will be zero. In contrast, if detections beyond RDR are considered (DDF case), the ITDP does not drop below 90% even when RT=10min, which we consider to be a very long reaction time. This is explained by the fact that animals can be detected further out, hence are available for detection in the safe zone more frequently. For the remainder of this article we will only consider the DDF case, because of its superior ITDP. In every operational setting, DDF is the realistic detection case, and the RDR scenario is only relevant for academic purposes.

**Figure 3:**
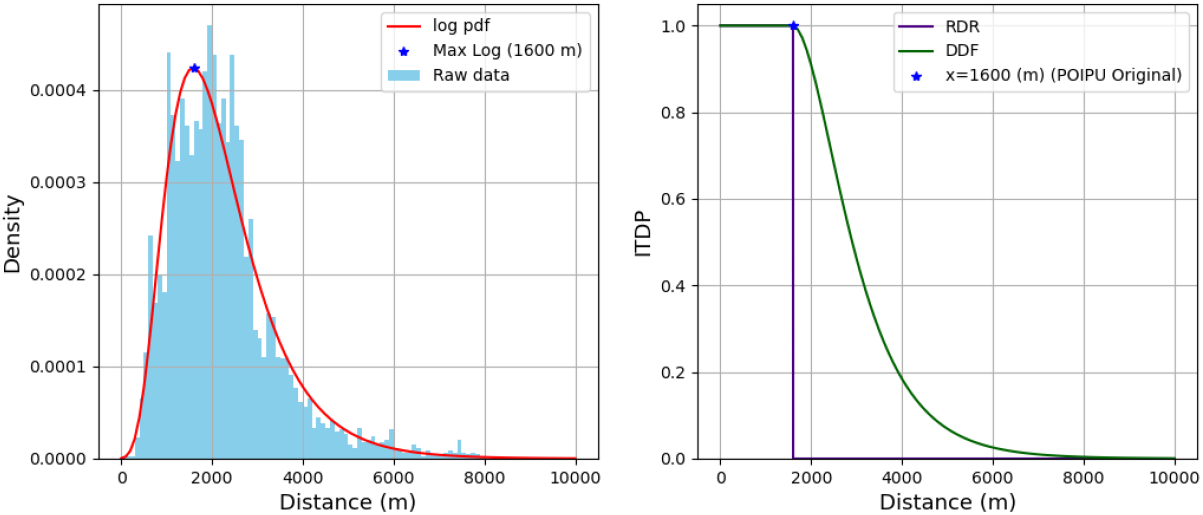
A) density distribution of detected blows, from humpback whales, as a function of distance. The data was collected in 2016 off the Poipu Shores, Hawaii, using a thermal IR imaging camera located 16m from the MSL. A log-normal function (log pdf, red) was fitted to the raw distribution. The maximum value reached by the log pdf is marked with a blue star (max log) at 1600m and depicts the Reliable Detection Range (RDR). B illustrates the RDR scenario (purple), where the probability of detection is equal to 1 for distances below or equal to the RDR, and the Data-driven Detection Function (DDF, green) where the function logarithmically decreases to 0 past the RDR.

**Figure 4:**
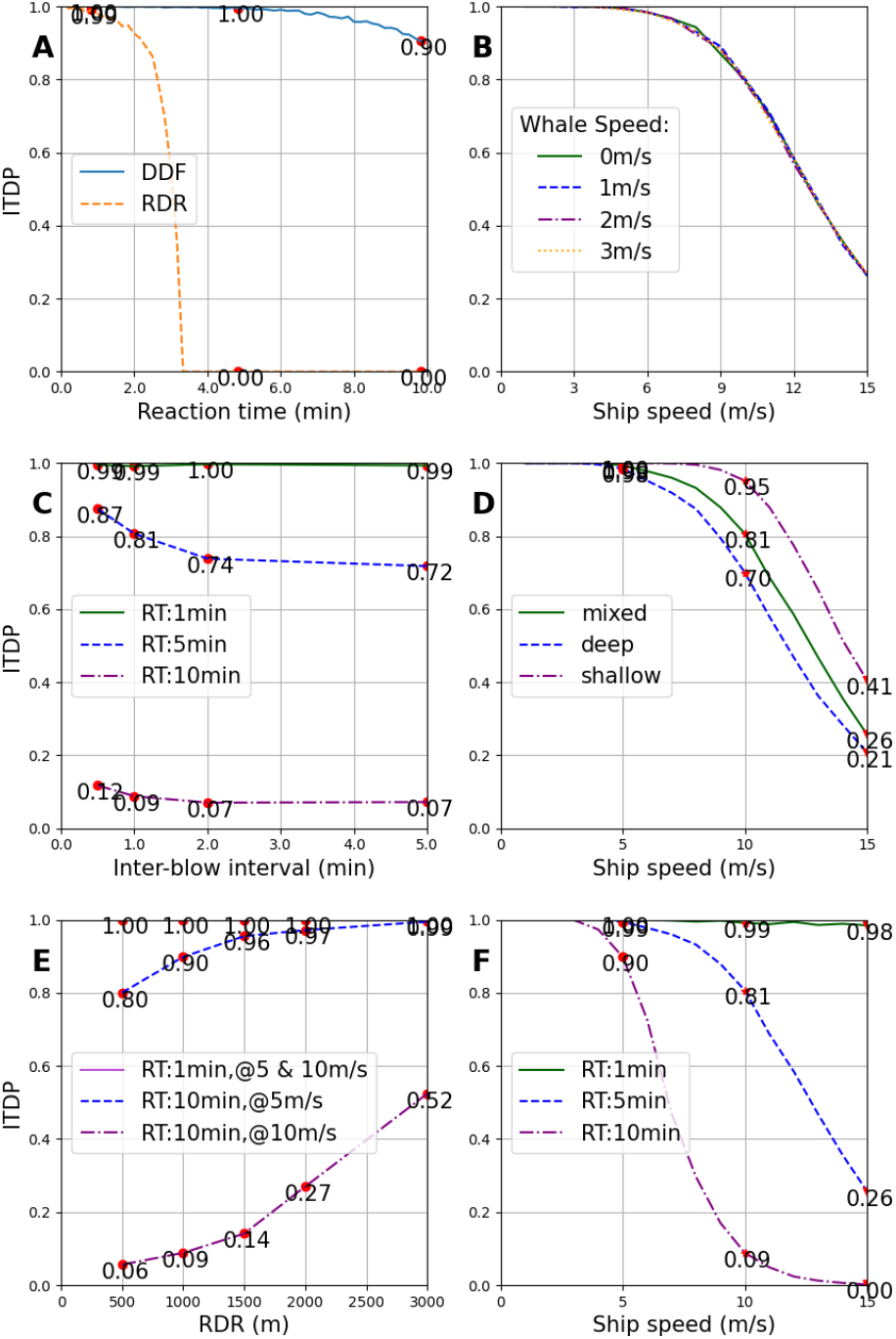
In-Time Detection Probability (ITDP) as a function of different parameters. A) Reaction Time (RT) for the two detection scenarios (RDR vs. DDF). B) Ship speeds for varying whale speeds (0,1,2,3m/s). C) Inter-Blow Intervals (IBI=30,60,120,300s) for different RTs (RT=1,5,10min). D) whale dive profile behavior (mixed,shallow,deep) as a function of ship speeds. E) Reliable Detection Ranges (RDRs=[500-3000m]) for varying RTs (RT=1,10min) and vessel speeds (5,10m/s). F) Reaction Times (RT=1,5,10min) as a function of ship speeds. All scenarios used DDF, RDR=1000m, DDF, IBI=60s, a mixed diving behavior, RT=300s if not otherwise mentioned.

### 3.2 Whale Speed

We find that in our model, the swimming speed of the animal has no effect on the ITDP (Figure 4B). This can be explained by the speed difference between the animal and the vessel. At slow vessel speeds, the safe zone is much larger than the danger zone, thus the animal is often available for detection regardless of its speed, hence we obtain high ITDP values. At higher vessel speeds, the ITDP drops. The decrease of the ITDP is independent from the whales’ speed and is only caused by increasing vessel speeds which drive up the danger zone range (DZR=RT*vessel speed). The only case where the whale’s speed could have an impact on ITDP is when an animal would be swimming (in a diving state) from outside the safe zone’s swath directly into the danger zone. The combinations of whale and vessel speed we chose apparently make this case impossible. Whales would need to be swimming at equal or higher speeds than the vessel, which is unrealistic for a prolonged duration. A comprehensive plot summarizing the ITDP results as a function of ship speeds for different whale speeds can be found in Appendix B.

### 3.3 Inter-Blow Interval (IBI)

Since inter-blow-interval data was not readily available, we simulated a range of possible IBIs (30, 60, 120, and 300sec) where each blow is detectable for 3 seconds. We find that the IBI does not reduce the ITPD by more than 16% (vessel speed=10m/s, RDR=1000m). In this case, we chose a rather high vessel speed of 10m/s compared to the other results presented, which were evaluated at 5m/s vessel speed, because at 5m/s the impact of the IBI is negligible. Furthermore, IBI only has an impact for RT larger than 300 sec. Overall, the impact of the IBI can be considered rather small.

### 3.4 Diving Behavior

Diving behavior of NARWs changes significantly throughout the year, depending on either the food availability at different depths of the ocean or the behavioral state of the animal (breeding vs foraging) (Murison & Gaskin 1989). We find that at low vessel speeds (≤5m/s), the behavior of the whale has negligible impact on the ITDP (98-100%). The diving behavior of the whale starts to have an impact for higher vessel speeds. At 10m/s, there is a 25% decrease of ITDP between shallow and deep dives. This observation can be explained by the fact that if the whale is on a deep dive, it is less often available for detection, i.e. fewer surfacings reduce the chances of detection in the safe zone, making the animal more vulnerable to strikes. The ITDP for mixed diving behavior lies as expected between deep and shallow diving behaviors which were used to compose it. To best generalize, we chose mixed diving behavior for the other presented results.

### 3.5 Reliable Detection Range (RDR)

The reliable detection range (RDR) is one of the main factors that determines the size of the safe zone. At a given vessel speed and reaction time (RT), the danger zone is constant in size, hence with increasing RDR, only the safe zone increases. As expected, with increasing RDR (i.e increasing safe zone), ITDP values increase as well. Because the size of the danger zone is determined by the vessel’s speed and RT, both parameters have to be considered jointly. For shorter RTs (RT=1min,5min), the ITDP [99-100%] is independent of both the vessel speeds (5 m/s vs. 10m/s) and the RDR [500-3000m] (Figure 4E). However, for longer RTs (RT=10min), higher vessel speeds negatively impact the ITDP. Specifically, we observe that the ITDP decreases by up to 81% [47-81%] when comparing results obtained at vessel speeds of 5m/s vs. 10m/s for RT=10min.

### 3.6 Vessel Reaction Time (RT)

We find that the impact of RT on the ITDP is highly dependent on the vessel’s speed. For short RTs, the ITDP is not significantly impacted by increasing vessel speeds (at most a 2% decrease across vessel speeds [1-15m/s] when RT=1min). On the other hand, longer RTs are very sensitive to higher vessel speeds with a decrease of up to 90% when comparing the ITDP of RT=10min at vessel speeds of 5m/s vs. 15m/s. At low vessel speeds (≤5m/s), the impact of RT is at most a 10% decrease (RT=1 min vs RT=10 min), compared to 90% for high vessel speeds (10m/s) (Figure 4F). This can be explained by higher RTs leading to larger danger zones, which increase the probability of a detectable whale surfacing in that zone rather than in the safe zone. A comprehensive plot summarizing the ITDP results as a function of ship speeds for all parameter modelled combinations can be found in Appendix C.

## 4 DISCUSSION

### 4.1 Lack of effective measures

Most of the current mitigation measures have significant limitations that impact their ability to properly and consistently protect NARWs and other marine mammals from vessel strike. Re-routing measures are not always feasible or when in place, rely on the cooperation of mariners. Vanderlaan & Taggart (2009) found that 71% of boats complied with the voluntary Roseway Basin ATBA in Canada, which led to a 82% decrease in lethal strikes to right whales in that region (Vanderlaan & Taggart 2009). To improve compliance, increased enforcement might benefit certain areas, but cannot always be achieved due to limited resources and/or large areas (Schoeman et al. 2020). Additionally, re-routing measures might help protect one species, but put at higher risk another one (Redfern et al. 2013). Similarly, when vessel speed restrictions are complied with, they are effective in decreasing the rate of strikes as well as the severity of injury (Vanderlaan & Taggart 2009). Edbon et al. (2020) observed that lethal vessel strikes to Bryde’s whales (*Balaenoptera edeni brydei*) in the Hauraki Gulf, New Zealand have halved since the implementation of speed reduction from 13.2 kt to 10 kt (Ebdon et al. 2020). However, the effectiveness of this measure entirely depends on compliance from mariners, which studies have shown to be low (Silber, Adams & Bettridge 2012, McKenna et al. 2012). For instance, along the coast of California, a voluntary conservation program was implemented asking vessels to reduce their speed to 10kt or less when transiting through a 75 nm stretch of shipping lanes in the region. McKenna et al. (2012) observed that speeds were not at or below the recommended 10 knots, nor were daily average speeds reduced during the requested periods. Hence, similarly as re-routing measures, increased enforcement should improve effectiveness of speed reduction programs when resources and geography allow. More recent statistics show increasing cooperation from the mariners to follow the 10kn speed rule. In 2018-2019, the highest level (81%) of mariner compliance with the speed rule was observed (NOAA Fisheries, Office of Protected Resources 2020). In most Seasonal Management Areas, 25% to 83% of the vessels maintained a speed under 10 kn (NOAA Fisheries, Office of Protected Resources 2020). Low numbers can be explained by the economic impact resulting from longer transits. NOAA Fisheries estimates it to be around $28.3 to $38.4 million annually, with the majority of the cost (50-70%) falling on the container ship sector (NOAA Fisheries, Office of Protected Resources 2020). The addition of vessel-based mitigation measures might be beneficial to improve effectiveness and compliance of current mitigation strategies. The purpose of our model is to assess in which scenarios, vessel-based whale strike mitigation would be useful to enhance protection of large whales. To this end, we have to consider the impact it would have (e.g. the ITDP) in different real-world scenarios.

### 4.2 Vessel Parameters Dependency

#### 4.2.1 Low vessel speed scenario

The most important finding is that in slow-speed environments, such as speed restricted zones, vessel-based whale detection systems for strike mitigation would provide a high protection for the animals. Specifically, if a vessel travels slowly (5m/s), any form of detection system with at least 1000m of reliable detection range will lead to high in-time detection probabilities (>90%). Detection ranges of 1000m can be achieved on the majority of vessels that provide ¿5m elevation (Zitterbart et al. 2020). Due to the slow vessel-speed, parameters such as the reaction time of the vessel become less relevant. The reaction time as we use it is mainly determined by the vessel’s maneuverability (which cannot be changed) and the time between detection and alert. The fact that the reaction time is of less relevance during slower travel is a key finding as it opens up the possibility for remote validation of vessel-based detections. One could imagine a system where automatic detections are transmitted to a data-center in near-real time and validated immediately. Turn-around times would likely be on the order of minutes, short enough to alert the vessel about the whale’s presence without any false alerts. We consider false alerts to be a major reason why automatic vessel-based whale detection systems could not be directly used by the vessel’s crew. We speculate that even at relatively low false alert rates (6 per hour, (Zitterbart et al. 2013)), a vessel’s crew is likely to soon ignore warnings.

### 4.2.2 High vessel speed scenario

In areas where vessel speed is not regulated and vessels travel at higher speeds (5-15m/s), we can group vessels into three classes. When vessels have a high maneuverability and have the capability to change velocity quickly, vessel-based whale detection systems can be very effective (ITDP > 98%), under the conditions that the reliable detection range is at least 1km and that the time from detection to alert is minimal (on the order of seconds). Currently, such conditions could only be met with dedicated observers on-board, who could either be scanning the ocean surface, or validate automatic detections provided by a whale detection system. Obtaining such a quick response comes with great efforts and costs, and hence is unlikely to be implemented on a large scale. For vessels with longer reaction times (lower maneuverability and/or longer detection-to-alert time), a whale detection system can still be effective if the reliable detection range is increased to several kilometers (+58% RDR=3000m vs. RDR=1000m when RT=5min and vessel speed=15m/s). This is valid for a large group of vessels (e.g. cruise vessels) that have high enough elevations to provide large detection ranges for the whale detection system (Zitterbart et al. 2020). Thus, longer reaction times can be off-setted by more advanced detection systems with larger reliable detection ranges or by slowing down the vessel’s speed. However, for vessels that travel at very high speeds, have very poor maneuverability, and might travel in shipping lanes, where quick maneuvers are not feasible (e.g. supertankers, large container vessels), no currently available vessel-based detection methods would provide enough detection range for effective protection.

### 4.3 Whale parameters dependency

Another finding of this simulation study is that animal behavioral factors such as swimming speeds, diving profiles and the inter-blow interval are not as impactful on the detection as the vessels’ parameters. Our simulation showed that varying whale speeds does not have any impact on the ITDP. This can be explained by the fact that NARWs are slow swimmers (∼ 0.36m/s on average) (Hain et al. 2013). Hence, at such low swimming speeds, their movement becomes negligible compared to the faster vessel speeds. Thus, we learn that in the case of NARWs, the animals’ horizontal movements do not need to be considered for the design of vessel-based detection systems (e.g. field of view), which can have a significant impact on the system costs, and potentially negatively impact their wide-spread use. Our study confirmed that NARW going on deep dives are more prone to collisions as they have fewer surfacings, compared to surface feeding NARWs (25% decrease at 10m/s vessel speed). However, our study also highlights that NARW’s behavior is only impactful when vessels are traveling at high speeds (10+m/s). Slower vessels have an equally high ITDP (98+%) regardless of the NARW’s diving behavior (Figure 4D). Hence, a large-scale mitigation measure, such as a speed restriction zone, would help alleviate the impact of whales’ diving behavior, and make the use of vessel-based mitigation systems most effective.

## 5 CONCLUSIONS

To summarize, under the right conditions (slow average vessel speed, high maneu-verability) vessel-based whale detection systems can be very effective for whale-strike mitigation and a large-scale deployment of such systems in high-risk areas could effectively reduce whale-strikes. While certain vessel classes cannot directly benefit from a vessel-based whale detection system, the information obtained and shared from other vessels equipped with such a system, could help estimate a near real-time distribution of whales in critical areas and improve the large-scale dynamic management efforts. With future technological improvements on the hardware (larger detection ranges) and software (fewer false positives), vessel strike mitigation systems could become a standard tool in the maritime industry.

## Supporting information

Case parameters

## 6 ACKNOWLEDGMENT

This work was supported by the Navy’s Office of Naval Research (ONR) under award number N000141912669. Additionally, we would like to thank Mark Baum-gartner for providing North Atlantic Right Whales dive profile data collected in 2000 and 2001.

## A Appendix

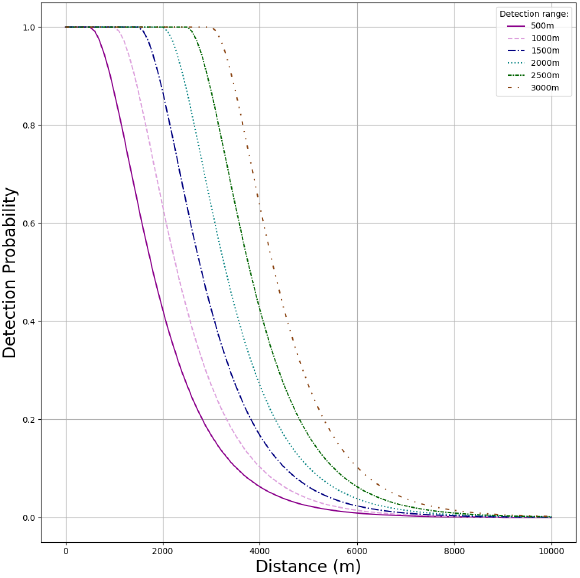

Detection Function as a function of distance (m) from the ship and detection range (m)

## B Appendix

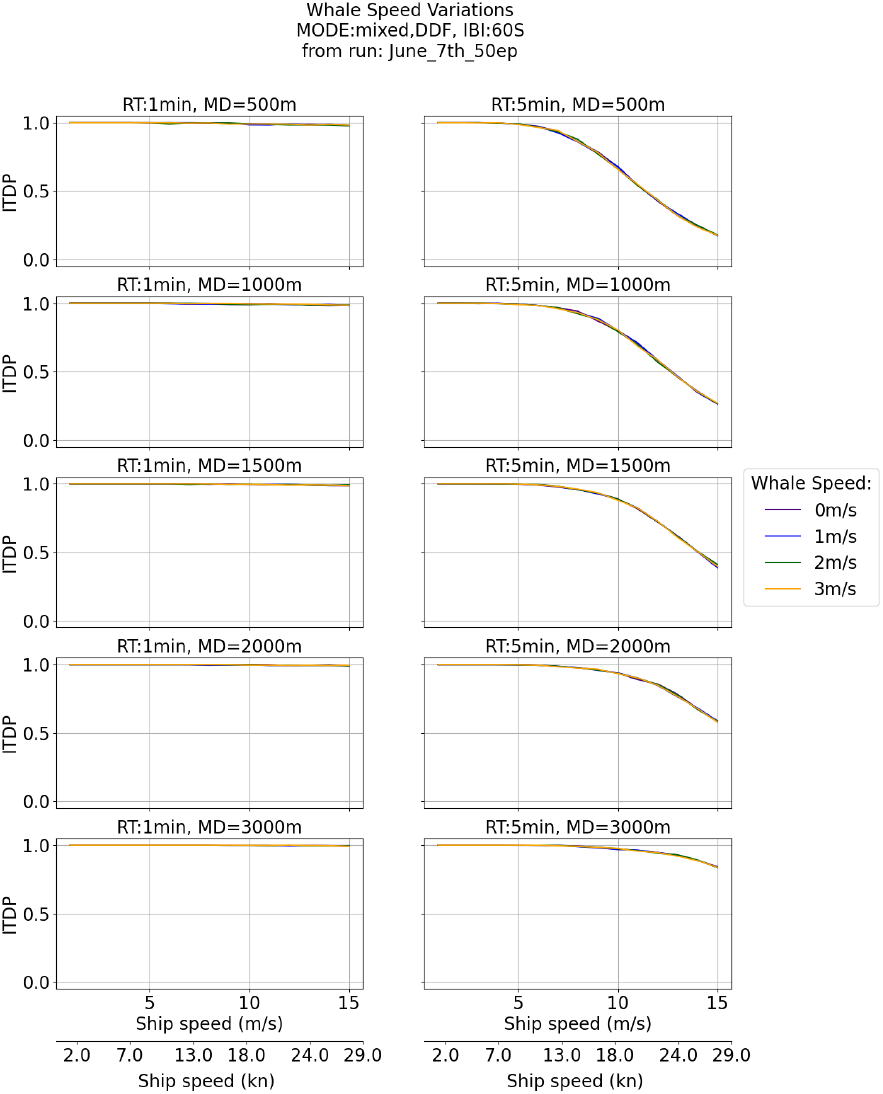

Comprehensive results of ITDP as a function of ship speeds (m/s) for varying whale speeds (m/s)

## C Appendix

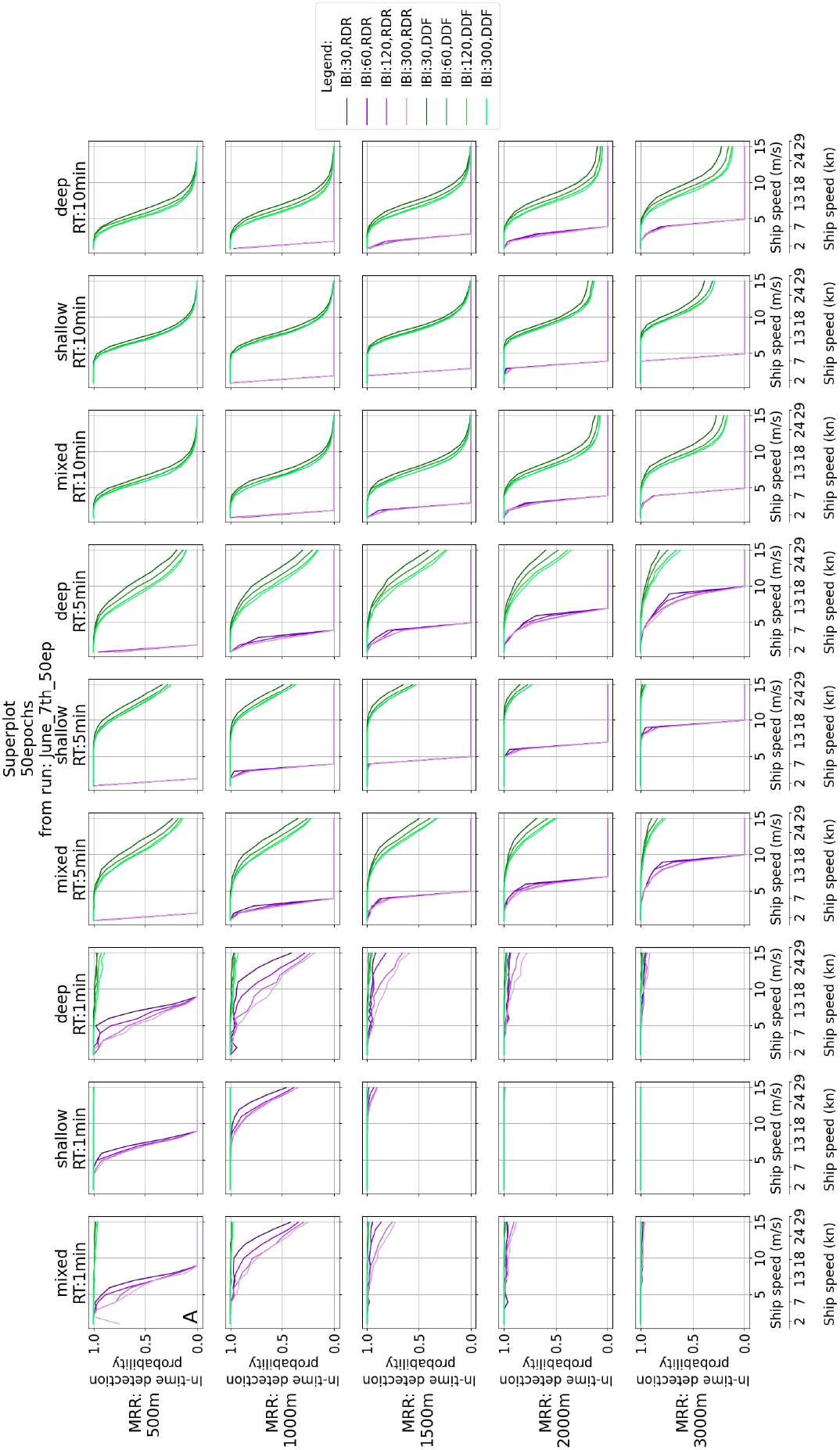

Comprehensive results of ITDP for varying ship speed (m/s) for all modelled parameters combination

## LITERATURE CITED

André, M., Houégnigan, L., van der Schaar, M., Delory, E., Zaugg, S., Sánchez, A. M. & Mas, A. (2011), Localising Cetacean Sounds for the Real-Time Mitigation and Long-Term Acoustic Monitoring of Noise, IntechOpen.

Baumgartner, M. F., Bonnell, J., Parijs, S. M. V., Corkeron, P. J., Hotchkin, C., Ball, K., Pelletier, L.-P., Partan, J., Peters, D., Kemp, J., Pietro, J., Newhall, K., Stokes, A., Cole, T. V. N., Quintana, E. & Kraus, S. D. (2019), ‘Persistent near real-time passive acoustic monitoring for baleen whales from a moored buoy: System description and evaluation’, Methods in Ecology and Evolution 10(9), 1476–1489.

Baumgartner, M. & Mate, B. (2003), ‘Summertime foraging ecology of North Atlantic right whales’, Marine Ecology Progress Series 264, 123–135.

Baumgartner, M., Wenzel, F., Lysiak, N. & Patrician, M. (2017), ‘North Atlantic right whale foraging ecology and its role in human-caused mortality’, Marine Ecology Progress Series 581, 165–181.

Campbell-Malone, R., Barco, S. G., Daoust, P.-Y., Knowlton, A. R., McLellan, W. A., Rotstein, D. S. & Moore, M. J. (2008), ‘Gross and Histologic Evidence of Sharp and Blunt Trauma in North Atlantic Right Whales (Eubalaena glacialis) Killed by Vessels’, Journal of Zoo and Wildlife Medicine 39(1), 37–55.

Conn, P. & Silber, G. (2013), ‘Vessel speed restrictions reduce risk of collision-related mortality for North Atlantic right whales’, Ecosphere 4, art43.

Ebdon, P., Riekkola, L. & Constantine, R. (2020), ‘Testing the efficacy of ship strike mitigation for whales in the Hauraki Gulf, New Zealand’, Ocean & Coastal Management 184, 105034.

Fisheries, N. (Thu, 06/03/2021 - 11:54), ‘North Atlantic Right Whale — NOAA Fisheries’, https://www.fisheries.noaa.gov/species/north-atlantic-right-whale.

Fisheries, N. (Thu, 06/03/2021 - 12:38), ‘Reducing Vessel Strikes to North Atlantic Right Whales — NOAA Fish-eries’, https://www.fisheries.noaa.gov/national/endangered-species-conservation/reducing-vessel-strikes-north-atlantic-right-whales.

Flynn, K. R. & Calambokidis, J. (2019), ‘Lessons From Placing an Observer on Commercial Cargo Ships Off the U.S. West Coast: Utility as an Observation Platform and Insight Into Ship Strike Vulnerability’, Frontiers in Marine Science 6.

Gende, S. M., Hendrix, A. N., Harris, K. R., Eichenlaub, B., Nielsen, J. & Pyare, S. (2011), ‘A Bayesian approach for understanding the role of ship speed in whale–ship encounters’, Ecological Applications 21(6), 2232–2240.

Hain, J. H. W., Hampp, J. D., McKenney, S. A., Albert, J. A. & Kenney, R. D. (2013), ‘Swim Speed, Behavior, and Movement of North Atlantic Right Whales (Eubalaena glacialis) in Coastal Waters of Northeastern Florida, USA’, PLOS ONE 8(1), e54340.

Hayes, S. A., Josephson, E., Cole, T. V., Engleby, L., Garrison, L. P., Hatch, J., Henry, A., Horstman, S. C., Litz, J., Lyssikatos, M. C., Mullin, K. D., Orphanides, C., Pace, R. M., Palka, D. L., Soldevilla, M. & Wenzel, F. W. (2018), US Atlantic and Gulf of Mexico Marine Mammal Stock Assessments - 2017: (second edition), Technical report, US DEPARTMENT OF COMMERCE, National Oceanic and Atmospheric Administration, National Marine Fisheries Service.

Johnson, H. D., Baumgartner, M. F. & Taggart, C. T. (2020), ‘Estimating North Atlantic right whale (Eubalaena glacialis) location uncertainty following visual or acoustic detection to inform dynamic management’, Conservation Science and Practice 2(10), e267.

Knowlton, C. (2020), ‘Real-time Passive Acoustic Sensors’, https://dosits.org/galleries/technology-gallery/observing-and-monitoring-marine-animals/real-time-passive-acoustic-sensors/.

Laist, D. W., Knowlton, A. R., Mead, J. G., Collet, A. S. & Podesta, M. (2001), ‘Collisions Between Ships and Whales’, Marine Mammal Science 17(1), 35–75.

McKenna, M. F., Katz, S. L., Condit, C. & Walbridge, S. (2012), ‘Response of Commercial Ships to a Voluntary Speed Reduction Measure: Are Voluntary Strategies Adequate for Mitigating Ship-Strike Risk?’, Coastal Management 40(6), 634–650.

Moore, M., Rowles, T., Fauquier, D., Baker, J., Biedron, I., Durban, J., Hamilton, P., Henry, A., Knowlton, A., McLellan, W., Miller, C., Pace, R., Pettis, H., Raverty, S., Rolland, R., Schick, R., Sharp, S., Smith, C., Thomas, L., van der Hoop, J. & Ziccardi, M. (2021), ‘REVIEW Assessing North Atlantic right whale health: Threats, and development of tools critical for conservation of the species’, Diseases of Aquatic Organisms 143, 205–226.

Murison, L. D. & Gaskin, D. E. (1989), ‘The distribution of right whales and Zooplankton in the Bay of Fundy, Canada’, Canadian Journal of Zoology 67(6), 1411–1420.

NATIONS, U. (n.d.), Review of Maritime Transport 2020, Technical report, UNITED NATIONS CONFERENCE ON TRADE AND DEVELOPEMENT.

NOAA Fisheries, Office of Protected Resources (2020), North Atlantic Right Whale (Eubalaena glacialis) Vessel Speed Rule Assessment, Technical report.

Pace, R. M., Williams, R., Kraus, S. D., Knowlton, A. R. & Pettis, H. M. (2021), ‘Cryptic mortality of North Atlantic right whales’, Conservation Science and Practice 3(2).

Parks, S. E., Warren, J. D., Stamieszkin, K., Mayo, C. A. & Wiley, D. (2012), ‘Dangerous dining: Surface foraging of North Atlantic right whales increases risk of vessel collisions’, Biology Letters 8(1), 57–60.

Peel, D., Smith, J. N. & Childerhouse, S. (2018), ‘Vessel Strike of Whales in Australia: The Challenges of Analysis of Historical Incident Data’, Frontiers in Marine Science 5.

Pyc, C., Geoffroy, M. & Knudsen, F. (2015), ‘An evaluation of active acoustic methods for detection of marine mammals in the Canadian Beaufort Sea’, Marine Mammal Science 32.

Redfern, J. V., McKenna, M. F., Moore, T. J., Calambokidis, J., Deangelis, M. L., Becker, E. A., Barlow, J., Forney, K. A., Fiedler, P. C. & Chivers, S. J. (2013), ‘Assessing the risk of ships striking large whales in marine spatial planning’, Conservation Biology: The Journal of the Society for Conservation Biology 27(2), 292–302.

Schoeman, R. P., Patterson-Abrolat, C. & Plön, S. (2020), ‘A Global Review of Vessel Collisions With Marine Animals’, Frontiers in Marine Science 7.

Sébe, M., Kontovas, C. A. & Pendleton, L. (2019), ‘A decision-making framework to reduce the risk of collisions between ships and whales’, Marine Policy 109, 103697.

Services, N. F. (n.d.), ‘Compliance Guide for Right Whale Ship Strike Reduction Rule’, p. 2.

Ships’ Routeing (n.d.), https://www.imo.org/en/OurWork/Safety/Pages/ShipsRouteing.aspx.

Silber, G., Adams, J. & Bettridge, S. (2012), ‘Vessel operator response to a voluntary measure for reducing collisions with whales’, Endangered Species Research 17, 245–254.

Silber, G. K., Vanderlaan, A. S., Tejedor Arceredillo, A., Johnson, L., Taggart, C. T., Brown, M. W., Bettridge, S. & Sagarminaga, R. (2012), ‘The role of the International Maritime Organization in reducing vessel threat to whales: Process, options, action and effectiveness’, Marine Policy 36(6), 1221–1233.

van der Hoop, J. M., Vanderlaan, A. S. M., Cole, T. V. N., Henry, A. G., Hall, L., Mase-Guthrie, B., Wimmer, T. & Moore, M. J. (2015), ‘Vessel Strikes to Large Whales Before and After the 2008 Ship Strike Rule’, Conservation Letters 8(1), 24–32.

Vanderlaan, A. S. M. & Taggart, C. T. (2007), ‘Vessel Collisions with Whales: The Probability of Lethal Injury Based on Vessel Speed’, Marine Mammal Science 23(1), 144–156.

Vanderlaan, A. S. M. & Taggart, C. T. (2009), ‘Efficacy of a Voluntary Area to Be Avoided to Reduce Risk of Lethal Vessel Strikes to Endangered Whales’, Conservation Biology 23(6), 1467–1474.

Verfuss, U., Gillespie, D., Gordon, J., Marques, T., Miller, B., Plunkett, R., Theriault, J., Tollit, D., Zitterbart, D., Hubert, P. & Thomas, L. (2018), ‘Comparing methods suitable for monitoring marine mammals in low visibility conditions during seismic surveys’, Marine Pollution Bulletin 126, 1–18.

Weinrich, M., Pekarcik, C. & Tackaberry, J. (2010), ‘The effectiveness of dedicated observers in reducing risks of marine mammal collisions with ferries: A test of the technique’, Marine Mammal Science 26(2), 460–470.

Wikipedia (2021), ‘Container ship’, Wikipedia Foundation.

Wiley, D. N., Mayo, C. A., Maloney, E. M. & Moore, M. J. (2016), ‘Vessel strike mitigation lessons from direct observations involving two collisions between non-commercial vessels and North Atlantic right whales (Eubalaena glacialis)’, Marine Mammal Science 32(4), 1501–1509.

Zitterbart, D. P., Kindermann, L., Burkhardt, E. & Boebel, O. (2013), ‘Automatic Round-the-Clock Detection of Whales for Mitigation from Underwater Noise Impacts’, PLOS ONE 8(8), e71217.

Zitterbart, D. P., Smith, H. R., Flau, M., Richter, S., Burkhardt, E., Beland, J., Bennett, L., Cammareri, A., Davis, A., Holst, M., Lanfredi, C., Michel, H., Noad, M., Owen, K., Pacini, A. & Boebel, O. (2020), ‘Scaling the Laws of Thermal Imaging–Based Whale Detection’, Journal of Atmospheric and Oceanic Technology 37(5), 807–824.

